# Synthetically mannosylated antigens induce antigen-specific humoral tolerance and reduce anti-drug antibody responses to immunogenic biologics

**DOI:** 10.1101/2023.04.07.534593

**Authors:** Rachel P. Wallace, Kirsten C. Refvik, Jennifer T. Antane, Kym Brünggel, Andrew C. Tremain, Michal R. Raczy, Aaron T. Alpar, Mindy Nguyen, Ani Solanki, Anna J. Slezak, Elyse A. Watkins, Abigail L. Lauterbach, Shijie Cao, D. Scott Wilson, Jeffrey A. Hubbell

## Abstract

Immunogenic biologics trigger an anti-drug antibody (ADA) response in patients, which reduces efficacy and increases adverse reactions. Our laboratory has previously shown that targeting protein antigen to the liver microenvironment can reduce antigen-specific T cell responses; herein, we present a strategy to increase delivery of otherwise immunogenic biologics to the liver via conjugation to a synthetic mannose polymer (p(Man)). This delivery leads to reduced antigen-specific T follicular helper cell and B cell responses resulting in diminished ADA production, which is maintained throughout subsequent administrations of the native biologic. We found that p(Man)-antigen treatment impairs the ADA response against recombinant uricase, a highly immunogenic biologic, without a dependence on hapten immunodominance or control by Tregs. We identify increased TCR signaling and increased apoptosis and exhaustion in T cells as effects of p(Man)-antigen treatment via transcriptomic analyses. This modular platform may enhance tolerance to biologics, enabling long-term solutions for an ever-increasing healthcare problem.

## Introduction

Since the approval of recombinant insulin in 1982, the development of therapeutic proteins, or biologics, has exploded, such that in 2020 biologics accounted for 20% of FDA approvals, a number that is expected to increase in the years to come^1^. Novel therapeutic classes such as monoclonal antibodies and enzyme replacement therapies enabled increasingly precise ways to treat or cure debilitating genetic diseases, cancer and autoimmunity. Compared to traditional small-molecule therapeutics, the protein composition of biologics enable them to directly interact with native pathways and mimic endogenous molecules to improve efficacy, but it also makes them vulnerable to recognition by the immune system. The immune response is primarily due to a T cell-dependent antibody response against the protein drug^2,3^. At sufficient levels, these anti-drug antibodies (ADAs) generated in response to the therapeutic can cause a large number of complications including neutralization of the drug’s therapeutic function, premature drug clearance, hypersensitivity and life-threatening anaphylaxis reactions upon repeated administration^4–8^.

A biologic’s immunogenicity is determined by a variety of factors including characteristics of the protein such as its homology to native human proteins expressed in that individual, the number of subunits, and post-translational modifications, as well as genetic factors and dose route, quantity, and frequency^3^. The pharmaceutical industry has employed a number of strategies to try to reduce immunogenicity throughout the development of the biologic including selecting sequences with maximum homology to human proteins, however immunogenicity remains a hurdle for regulatory approval^9,10^. Strategies to design less immunogenic proteins using MHC associated peptide proteomics, predictive algorithms and machine learning^11–16^ are under development, but the only treatments currently available to patients who require immunogenic drugs is to co-administer broad immunosuppressants such as methotrexate and thiopurines, which leave patients at increased risk for infection and malignancy^17–21^. There is a critical need for an antigen-specific approach to reduce immunogenicity to biologics. Here we present a modular platform that targets protein therapeutics to the liver’s innate tolerance inducing environment, and thus reduces the immunogenicity of protein therapeutics, allowing post-tolerization administration of the native therapeutic.

Our lab has previously developed a liver-targeted amine-reactive glycoconjugate system and demonstrated that antigen-glycopolymer conjugates elicit tolerogenic antigen-specific T cell signatures after both prophylactic and therapeutic administration^22,23^. The polymeric N-acetyl galactosamine-decorated glycopolymer (p(GalNAc)) used previously were designed to target hepatic antigen presenting cells expressing C-type lectins containing the QPD binding motif (i.e. hepatocytes, Kupffer cells, and liver sinusoidal endothelial cells). We demonstrated that intravenous administration of out antigen-p(GalNAc) resulted in antigen presentation to CD4^+^ and CD8^+^ T cells in the liver’s innate tolerogenic environment, resulting in CD4^+^ and CD8^+^ T cell deletion and anergy. Due to the ubiquitous presence of mannose on the surface of endogenous and foreign antigens from yeast, bacteria, and viruses, many immune and parenchymal cells have evolved to recognize mannosylated antigens for uptake and presentation upon binding to EPN motif-containing lectins^24,25^. We previously developed a polymer containing mannose residues along with TLR7 agonists as a vaccine platform and demonstrated that mannose polymer binding to C-type lectins was critical for antigen uptake and presentation to a wide variety of professional APCs such as DCs and macrophages^26–28^. Here, we sought to utilize a pure mannose glycopolymer to act as a ligand for a broader range of mannose-binding lectins and investigate its potential as a tolerogenic therapy to reduce both T cell and B cell-mediated ADA responses to immunogenic therapeutics. We show that after intravenous injection our p(Man) polymer construct localizes to the liver as other macromolecules do when injected intravenously. We find this localization to be significantly more efficient than unconjugated antigen likely due to the polymer’s interactions with liver-resident APCs through their mannose-binding C-type lectin receptors. This increased antigen localization in the liver appears to result in reduced antigen-specific T cell, B cell and antibody responses to a variety of immunogenic proteins. In addition, through depletion studies and transcriptional analysis, we demonstrate that the reduced ADA response generated by our platform likely occurs due to a reduced follicular helper T cell response and is independent of regulatory T cells or hapten immunodominance, i.e. any immune dominance of the conjugated polymer.

## Results

### Conjugation to p(Man) increases antigen uptake and prolongs presentation in the liver

We sought to utilize a novel glycopolymer as a ligand for a broader range of mannose-binding type II C-type lectins and investigate its potential as a tolerogenic therapy (**Figure 1A, Supplemental Figures 1-3**). Mannose-decorated polymers (p(Man)) were synthesized via reversible addition fragmentation chain-transfer polymerization (RAFT) from a mannose-decorated monomer, N-(*2*-*hydroxyethyl*)*methacrylamide* —a biologically inert comonomer—and an azide-terminated RAFT agent. The model antigen ovalbumin (OVA) was first modified with an azide- and amine-reactive di-functional polyethylene glycol linker that upon conjugation to the protein forms a self-immolative linkage that is relatively stable in the extracellular space, but degrades upon internalization, releasing the protein in its unmodified form^26^ (**Supplemental Figure 4A**). To measure the liver-targeting effects of p(Man) conjugation compared to the glycopolymers previously explored by our group, we performed a biodistribution study. Mice were treated with saline, dye-labeled OVA (OVA_647_), or OVA_647_ conjugated to p(Man), p(GalNAc), or a synthetic glycopolymer decorated with N-acetyl glucosamine (p(GlcNAc)) via intravenous injection (i.v.). Organs were collected 3 hr after injection and measured via IVIS. At the current dose, only the mice treated with p(Man)-OVA_647_ experienced a significant increase in liver fluorescence signal compared to the OVA_647_ control (**Figure 1B-C**). In addition, there was no off-target OVA_647_ signal detected in other organs (**Supplemental Figure 4B**).

**Figure 1:**
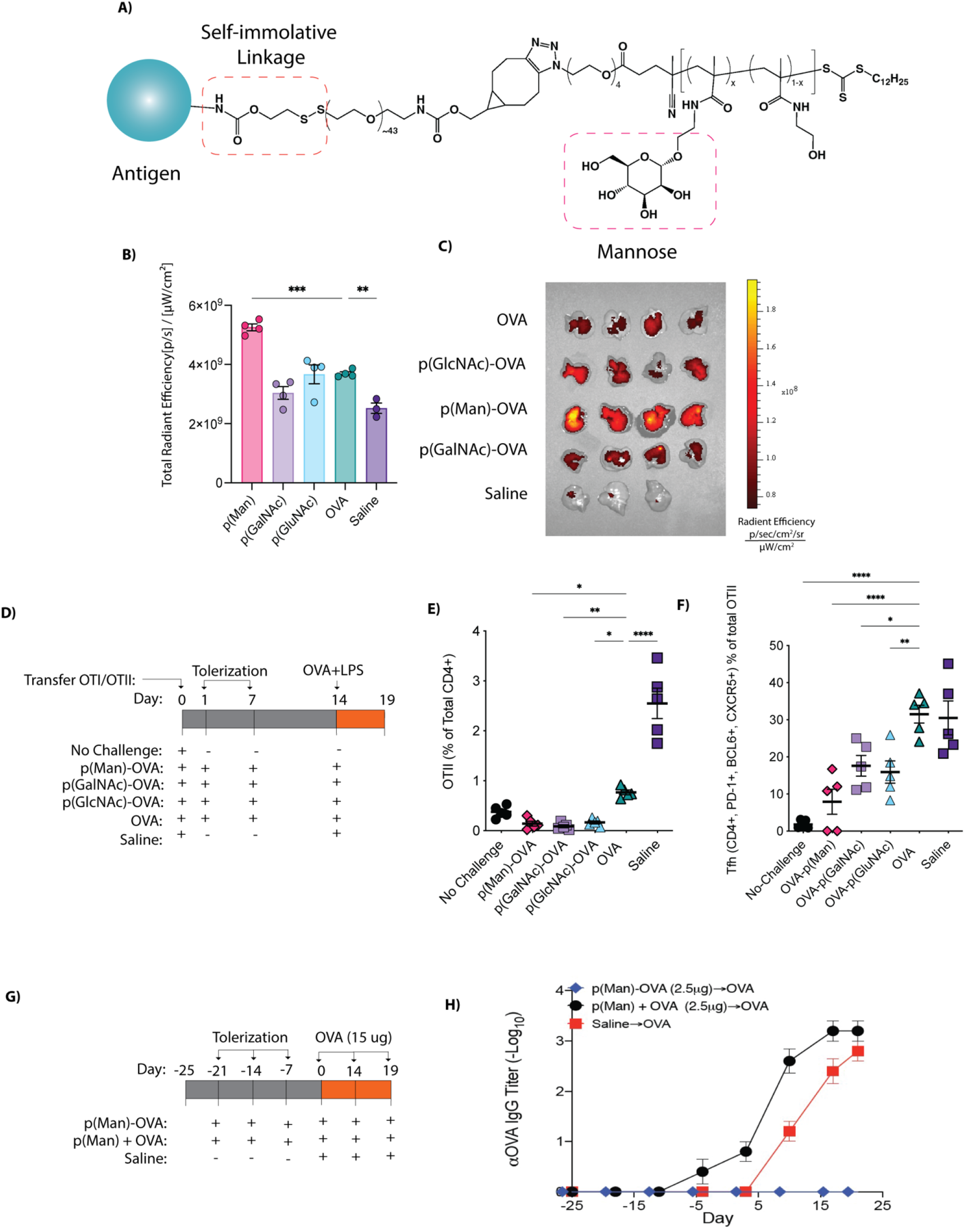
p(Man) conjugation increases delivery of antigen to the liver and impairs the activation of CD4^+^ antigen-specific T follicular helper cells. (A) Structure of the p(Man) polymer conjugated to protein antigen. (B) *N*=4 C57Bl/6 mice were administered saline, free OVA-AF647 or OVA-AF647 conjugated to glycopolymers p(Man), p(GlcNAc), and p(GalNAc). 3 hr after injection organs were isolated to measure radiant efficiency of the fluorescent antigen in the livers. (C) Representative IVIS images of the liver, treated as in (B). (D) 500,000 CD45.1 congenically labeled OT-I and OT-II T cells were adoptively transferred into *n*=5 C57Bl/6 mice. Mice were then treated with saline, free OVA, or OVA conjugated to p(Man), p(GalNAc) or p(GlcNAc) before vaccination with OVA+ LPS. OT-II cells were isolated on day 19 for phenotypic analysis. (E) Frequency of OT-II in the dLN after vaccination. (F) Frequency of PD-1^+^ Bcl6^+^ CXCR5^+^ T follicular helper (Tfh) cells within the OT-II population after vaccination. (G) Balb/c mice were treated weekly with p(Man)-OVA, a mixture of unconjugated p(Man) and OVA or saline for three weeks followed by a weekly challenge of high dose OVA. (H) Time course of OVA-specific IgG response represented as log_10_ titer. Symbols represent mean across *n*=8 mice. Data are shown as means ± SEM. Unless otherwise stated symbols represent individual mice and statistical differences in all graphs were determined by one-way ANOVA with Tukey’s *p<0.05, **p<0.01 and ***p<0.001.

### Pre-treatment with p(Man)-OVA results in decreased antigen-specific Tfh cells and prevents **α**OVA antibody response

We next investigated the effects of p(Man)-antigen therapy on antigen-specific T cell phenotypes using an OT-I/ OT-II vaccination model, in which adoptive transfer of OVA-specific CD8^+^ T cells (OT-I) or CD4^+^ T cells (OT-II) is followed by tolerization with p(Man)-antigen and then vaccination to challenge the tolerization regime (**Figure 1D**). When mice were given prophylactic treatment with p(Man)-OVA, we saw an impaired expansion of OVA-specific CD4^+^ T cells upon vaccination compared to the untreated mice (**Figure 1E**). Specifically, we found the greatest reduction in CD4^+^PD-1^hi^CD44^+^CXCR5^+^Bcl-6^+^ T follicular helper (Tfh) cells within the OT-II compartment after p(Man)-antigen treatment (**Figure 1F**). Given the essential role played by Tfh cells in germinal center formation, antibody affinity maturation, and memory B cell formation^29^, the reduction in Tfh cells prompted us to investigate the effects of p(Man)-antigen therapy on humoral immune responses. We treated mice weekly via i.v. injection with p(Man)-OVA, followed by weekly i.v. challenges of unmodified OVA (**Figure 1G**). We found that p(Man)-OVA antigen therapy prevented an antibody response to OVA, while treatment with p(Man) co-administered with unconjugated OVA expedited and increased the antibody response to OVA (**Figure 1H**).

### Pre-treatment with p(Man)-conjugated antigen prevents ADA responses to immunogenic therapeutic uricase

Once we established that p(Man)-OVA treatment can prevent ADA responses to OVA, we sought to investigate whether these tolerogenic effects would prevent a strong antibody response against a more immunogenic antigen like uricase. Uricase is an enzyme approved for the short-term treatment of hyperuricemia, however its long-term use is often limited by immunogenicity. Upon exposure to *Candida* uricase, mice generated a robust antigen-specific antibody response. This antibody response was shown to be T cell dependent and was completely abrogated when αCD4 and αCD8 blocking antibodies were administered (**Supplemental Figure 5A,B**).

We produced p(Man)-uricase conjugates and verified the reaction and the self-immolative characteristics of the linker via SDS-PAGE (**Supplemental Figure 6A**). While the enzymatic byproduct of uricase, hydrogen peroxide, is known to have an inflammatory influence on the immune system^30^, the enzymatic activity of uricase was measured and found to be significantly decreased upon conjugation to p(Man) and is unlikely to affect APCs upon uptake (**Supplemental Figure 6B,C**). We administered p(Man)-uricase, unmodified uricase or saline i.v. to mice weekly during a 3 wk tolerization window followed by weekly challenge doses of unmodified uricase (**Figure 2A**). Serum was collected weekly, and uricase-specific antibodies were measured via ELISA. The uricase-specific antibody response was represented as the area under the curve of the absorbance over background across a 10-fold log-transformed dilution series. Pre-treatment with p(Man)-uricase was found to significantly decrease the uricase-specific IgG response after high-dose antigen challenge compared to treatment with unmodified uricase and saline-treated mice (**Figure 2B,C**). This reduction in antibody response at endpoint was consistent across all subclasses of IgG (**Figure 2D**). To ensure this reduction in antibody response was consistent across immunogenic antigens, we repeated the experiment using *E. coli* asparaginase, an immunogenic protein used in oncology, and found consistent suppression of the antigen-specific IgG response after p(Man)-antigen treatment (**Supplemental Figure 7**).

**Figure 2:**
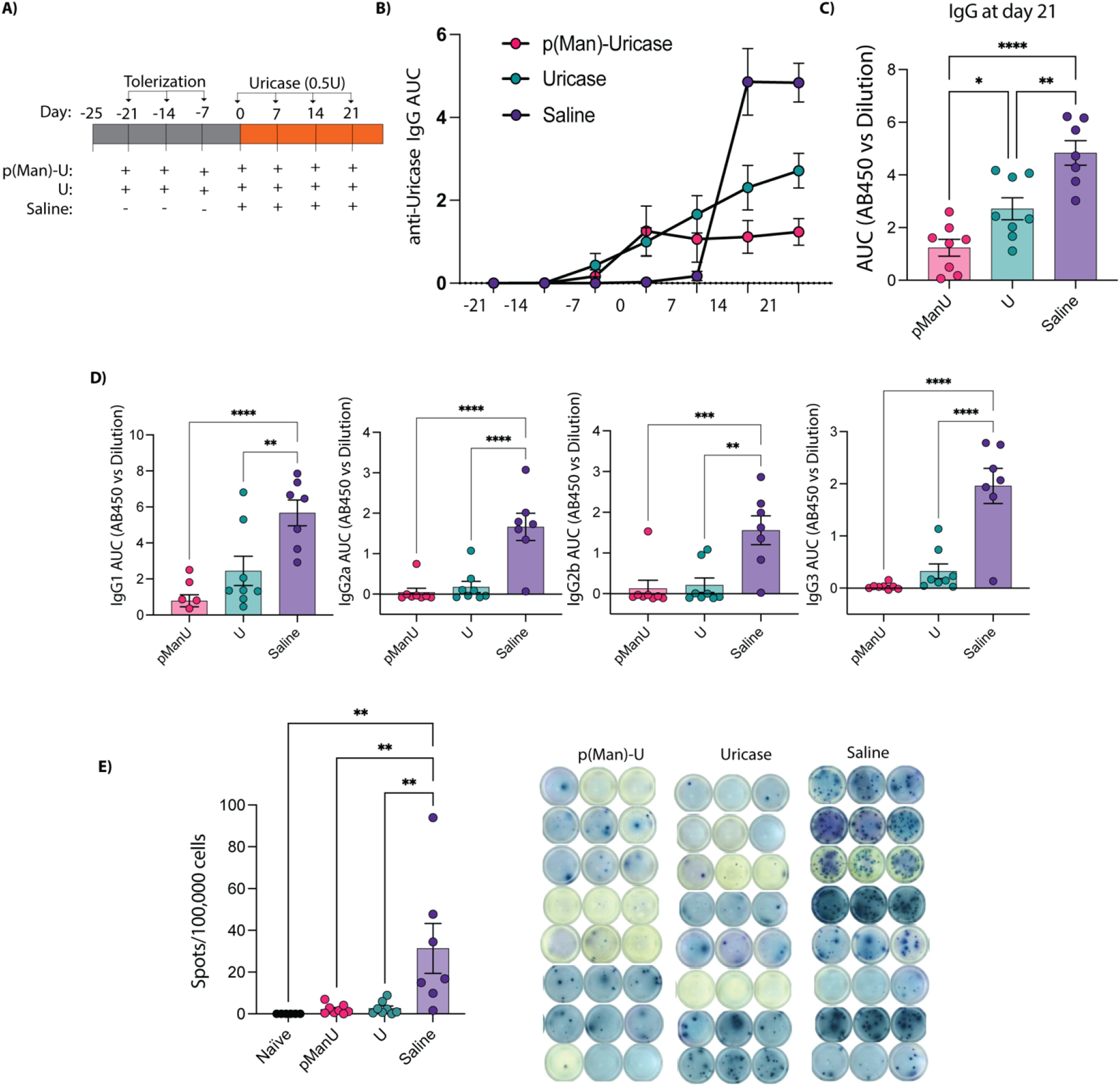
Prophylactic administration of p(Man)-antigen, impairs anti-drug antibody responses to immunogenic therapeutic uricase. (A) *N*=8 Balb/c mice were treated 3 times with p(Man)-uricase, free uricase or saline followed by a therapeutic dose of uricase weekly for 4 wk. (B) Time course of uricase-specific IgG response represented as the area under the curve of absorbance vs log-transformed dilution (AUC). Symbols represent mean across *n*=8 mice. (C) Uricase-specific IgG response at day 21 represented by AUC. (D) Comparison of uricase-specific IgG subclasses at day 21 represented by AUC. (E) Quantification and representative wells of the uricase-specific IgG-secreting splenocytes by ELISpot. Data are shown as means ± SEM. Unless otherwise stated symbols represent individual mice and statistical differences in all graphs were determined by one-way ANOVA with Tukey’s *p<0.05, **p<0.01 and ***p<0.001.

We next quantified the number of uricase-specific antibody-secreting cells using ELISpot. Only the untreated group was found to have significantly more uricase-specific antibody-secreting cells in the spleen on day 26 after initial uricase challenge, indicative of a smaller plasmablast and plasma cell response after both antigen and p(Man)-antigen pre-treatment (**Figure 2E**).

We next looked at the effect of p(Man)-uricase treatment on the B cell and T cell compartments (**Supplemental Figure 8A,B**). Fluorescent uricase-streptavidin multimers were used to identify antigen-specific B cells via flow cytometry (**Figure 3A**). Antigen-specificity was verified by incubating splenocytes from uricase-experienced mice with 5 mM uricase or OVA prior to multimer staining. Pre-incubation with uricase, but not OVA, resulted in the abrogation of uricase multimer binding (**Supplemental Figure 9**). Pre-treatment with p(Man)-uricase but not unmodified uricase prevented a significant rise in uricase-specific B cells in the spleen compared to the naïve controls as well as a noteworthy reduction in CD38^+^ IgD^-^ memory B cells and CD38^+^IgD^-^IgM^-^ class-switched memory B cells compared to the saline control treatment, which may indicate the potential for long-term B cell tolerance after p(Man)-antigen therapy (**Figure 3B**). We also saw trends towards reduction in plasmablasts and germinal center B cells, indicating a broadly reduced uricase-specific B cell response in the p(Man)-treated groups. The phenotype of the corresponding bulk B cell compartment, in contrast, showed no difference between treatment groups, indicating that p(Man)-uricase treatment is an antigen-specific treatment without broad non-specific effects on B cells (**Figure 3C**). Similarly, the cell counts of the bulk T cell compartment were consistent across groups, with the exception that all groups exposed to foreign uricase antigen had increased frequency of Tfh cells within the T cell compartment compared to the naïve control (**Supplemental Figure 10A,B**). The p(Man)-uricase treated group was the only group without a significant increase in overall cell numbers of Tfh cells and ICOS^hi^ Tfh cells compared to the naïve control (**Supplemental Figure 10B,C**).

**Figure 3:**
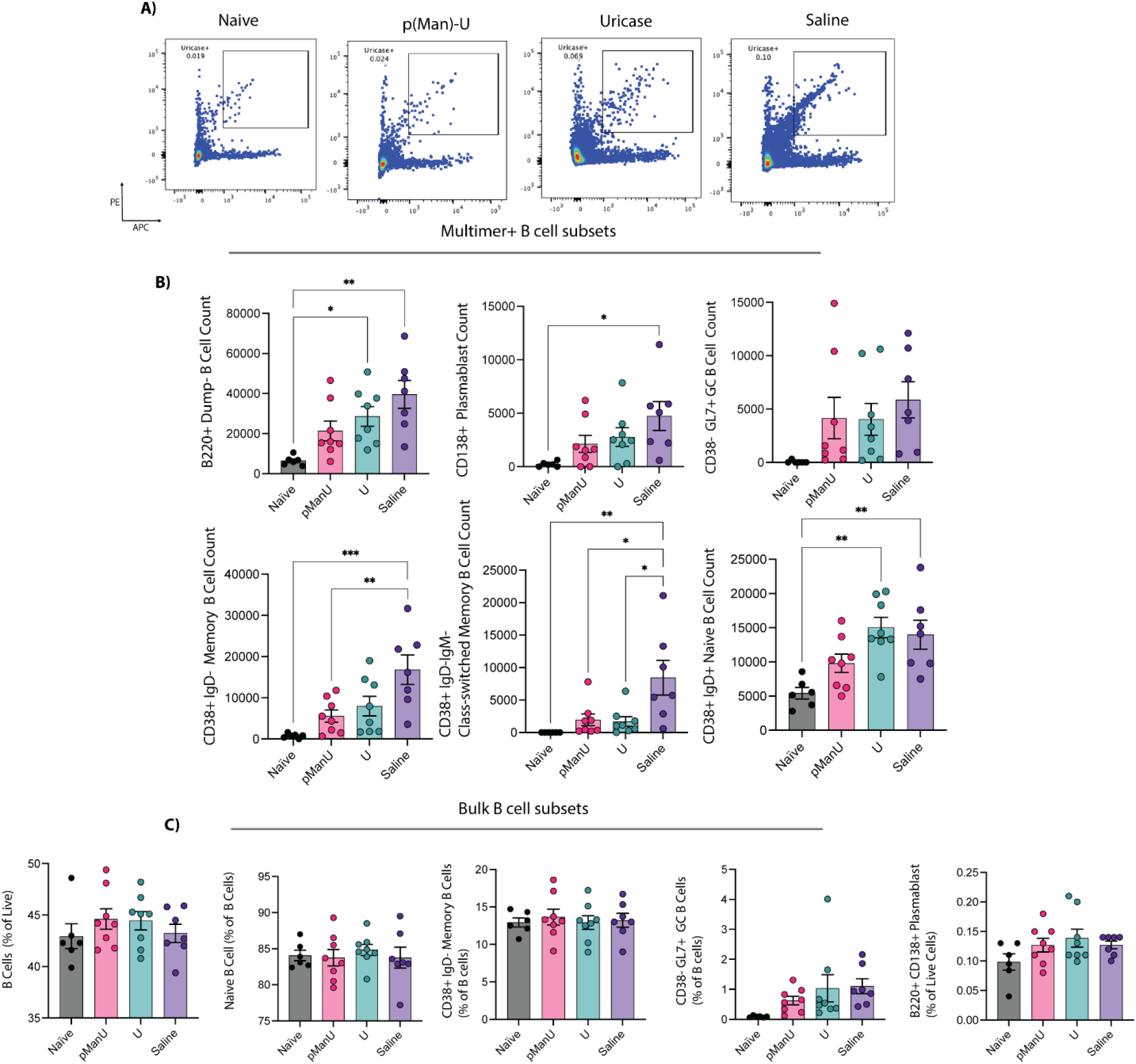
Antigen-specific memory B cell responses are impaired after prophylactic p(Man)-uricase treatment. (A) Representative flow plots of antigen-specific uricase-multimer double-positive B cells of naïve mice and uricase-challenged mice on day 25 after treatment with p(Man)-uricase, uricase, or saline. (B) Quantification of uricase-specific B220^+^ B cells, B220^+^CD138^+^ plasmablasts, B220^+^CD38^-^GL7^+^ germinal center B cells, CD38^+^IgD^-^ memory B cells, CD38^+^IgM^-^IgD^-^ class-switched memory B cells and CD38^+^IgD^+^ naïve B cells in the spleen on day 25 after treatment. (C) Quantification of bulk B220^+^ B cells as in (B). (D) Quantification of bulk CD3^+^ T cells, CD4^+^CXCR5^+^PD-1^+^Bcl6^+^ Tfh cells and ICOS^hi^ Tfh cells in the spleen on day 25 after treatment. Data are shown as means ± SEM. Unless otherwise stated symbols represent individual mice and statistical differences in all graphs were determined by one-way ANOVA with Tukey’s *p<0.05, **p<0.01 and ***p<0.001.

### Reduction in antigen-specific antibody response is not due to shift in the response to αp(Man) antibodies

After we established that pre-treatment with p(Man)-uricase leads to a reduced uricase-specific antibody response, we wanted to probe the potential mechanisms contributing to this effect. One possible mechanism leading to a reduced antigen-specific response is that the B cells recognize the p(Man) polymer instead of uricase and out-compete the uricase-specific B cells for T cell help in the germinal center, leading to a strong p(Man)-specific response instead of a uricase-specific response. To test the p(Man)-specific antibody response, we coated ELISA plates with uricase, p(Man)-uricase, p(Man)-OVA or p(Man) alone and measured antibodies from mice pre-treated with p(Man)-uricase, uricase or saline as in Figure 2A (**Figure 4A**). All of these mice were naïve to the OVA protein. We found the p(Man)-uricase treated mice generated no antibodies that bound p(Man)-OVA or p(Man) and the average antibody response to p(Man)-uricase was not higher than unmodified uricase in p(Man)-uricase treated mice (**Figure 4B**). These data indicate there was no antibody response to the p(Man) polymer regardless of treatment and thus that the possibility of immunodominance can be excluded.

**Figure 4:**
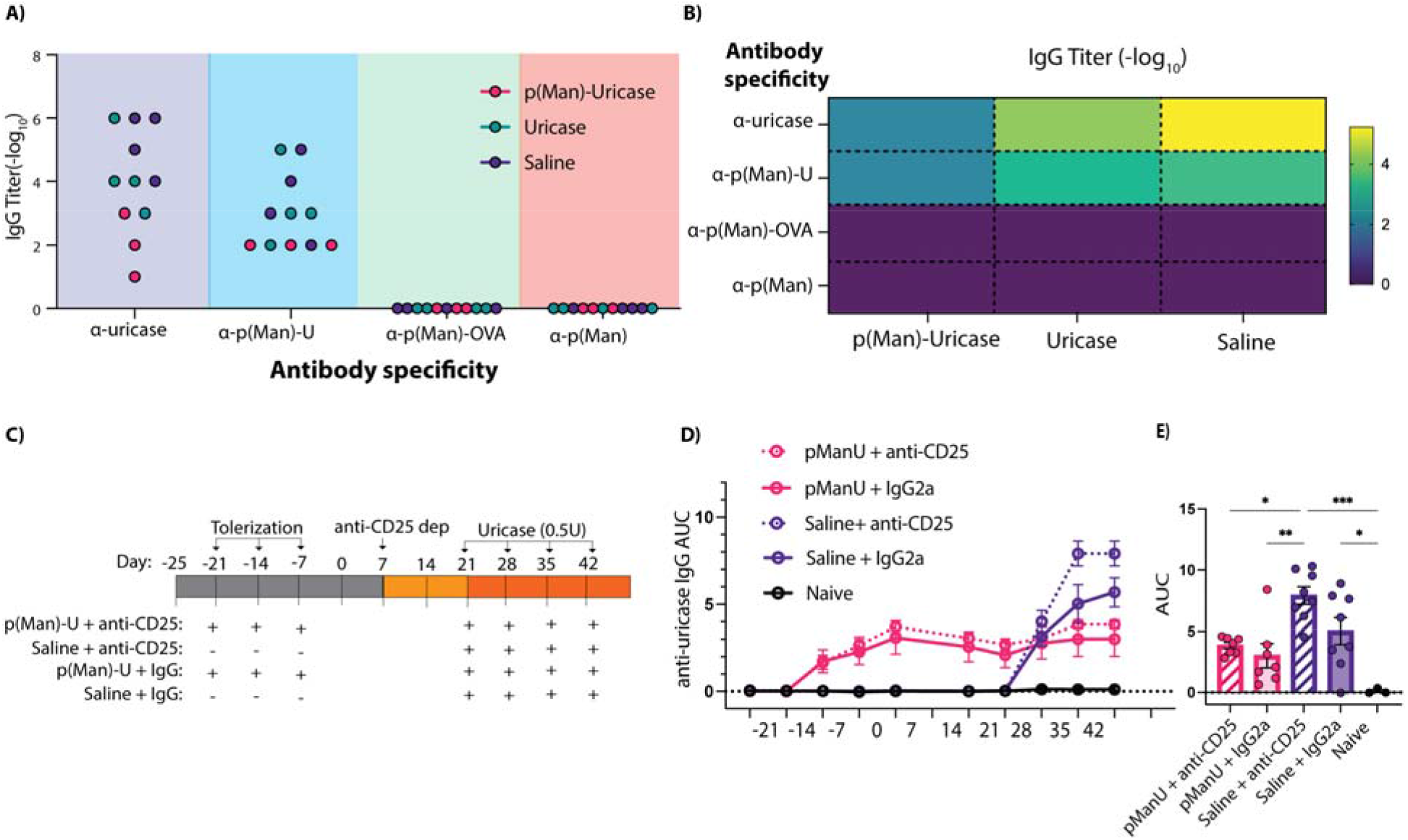
p(Man)-antigen treatment leads to an impaired antigen-specific antibody response independently of hapten immunodominance or regulatory T cell control. (A) Quantification of the antigen-specific antibody response to uricase, p(Man)-uricase, p(Man)-OVA or free p(Man) after treatment as in 2D represented as log_10_ titer. (B) Heatmap representation of the average log_10_ titer from each treatment group against each antigen as in (A). (C) N=8 Balb/c mice were treated three times with saline or p(Man)-uricase followed by administration of an isotype control or αCD25. High dose uricase injections were administered 2 wk after depletion for 4 wk. (D) Time course of the uricase-specific IgG response represented as AUC. Symbols represent mean across *n*=8 mice. (E) Uricase-specific IgG at day 45. Data are shown as means ± SEM. Unless otherwise stated symbols represent individual mice and statistical differences in all graphs were determined by one-way ANOVA with Tukey’s *p<0.05, **p<0.01 and ***p<0.001.

### Tregs are not required to maintain humoral tolerance generated by p(Man)-antigen treatment in an ADA context

We next decided to investigate if the antigen-specific T regulatory (Treg) cells induced by p(Man)-antigen treatment were required during high-dose challenge to maintain the reduced ADA response. In the OT-II vaccination study, we found that treatment with p(Man)-OVA resulted in an increased percentage of FoxP3^+^CD25^+^ Treg cells (**Supplemental Figure 11**). These cells could be responsible for the reduced antibody response through either Tfh suppression or direct B cell suppression. In order to probe the role of Tregs in the reduced uricase-specific antibody response, we administered αCD25 depleting antibody to the mice after tolerization and waited 2 wk for the antibody to be cleared before challenging with uricase (**Figure 4C**). We found that administration of the depleting antibody was not sufficient to prevent the reduction in antibody response seen after p(Man)-uricase treatment (**Figure 4D,E**). Therefore, active suppression of Tfh or B cells by Tregs during subsequent high dose-antigen challenge is likely not the mechanism by which p(Man)-antigen therapy imparts humoral immune tolerance in the ADA context.

### p(Man)-OVA treatment results in transcriptionally distinct profiles within OT-II cells

To further investigate the antigen-specific T cell-intrinsic mechanisms by which p(Man)-antigen therapy could result in reduced antigen-specific antibody responses, we performed bulk RNA sequencing on OT-II cells 5 days after antigen treatment (**Figure 5A**). We found the p(Man)-OVA treatment group resulted in 1,994 differentially expressed genes (DEGs) compared to the saline control, while treatment with unmodified OVA resulted in 132 DEGs compared to saline. Of the 1,994 DEGs altered by p(Man)-treatment, only 114 were found to also be differentially expressed in the OVA treatment comparison (**Figure 5B**). This indicates that within the antigen-specific CD4^+^ T cells, p(Man)-OVA treatment is driving transcriptional changes that are distinct from those up-regulated by native antigen presentation. Examination of the nature of the transcriptional distinctions between these two treatments provided evidence that, even at this early stage, pro-apoptotic and immunoregulatory markers distinguish the transcriptome after p(Man)-OVA treatment (**Figure 5C)**. While both treatments exhibit the anticipated activation and proliferative profiles of naïve T cells in the early stages after antigen recognition, overexpression of caspases and downregulation of heat shock proteins in the p(Man-OVA)-treated cells depicts a stronger pro-apoptotic profile, suggestive of a pro-tolerogenic deletion of these cells from the repertoire. Simultaneously, a stronger tolerogenic signature among the OT-II cells in the p(Man)-OVA group likely indicates the development of a persisting population of regulatory cells. Significant upregulation of co-inhibitory receptors, anergy-associated markers, and Treg-associated chemokine receptors like CCR10 all indicate that p(Man)-OVA treatment imprints a unique tolerogenic phenotype on these cells^31^ **(Figure 5D).** To understand which pathways these DEGs may be affecting in the p(Man)-OVA treatment compared to OVA treatment, we performed ingenuity pathway analysis (IPA). We identified 36 T cell-relevant immune pathways significantly altered in p(Man)-OVA treated groups compared to unmodified OVA (**Figure 5E**). The majority of these pathways were found to be associated with T cell activation and TCR signaling. Gene signatures were found for Th1, Th2, Tfh and Th17 pathways, although these pathways share many activation signatures such as increases in CD3ζ, NFAT, JAK and STAT signaling, which may explain why all are identified as significant. Some pathways associated with T cell activation such as Nur77 and TWEAK signaling are also associated with apoptosis and controlling T cell proliferation. The upregulation of signaling genes after p(Man)-OVA treatment is also significantly associated with an increase in cytokine signaling pathways such as IL-2, interferons and IL-23. Along with increased T cell signaling and activation, IPA identified pathways associated with exhaustion and apoptosis. The T cell exhaustion pathway is significantly upregulated due to increases in expression of co-inhibitory receptors CTLA4, EOMES, and PD-1. Upregulation of caspases and BAD are associated with apoptosis pathways, which are significantly upregulated in OT-II cells after p(Man)-OVA treatment. The combination of T cell activation along with upregulated apoptosis and exhaustion pathways suggest that p(Man)-OVA may lead to a deficient antigen-specific T follicular helper response through the activation of T cells towards different pathways such as Th1, Th2 or Th17, or alternatively that the increased antigen experience and activation signaling may lead to broad apoptosis and exhaustion of OVA-specific T cells. Alongside responses leading to lack of T cell help, we observe at this early time point of 5 days post p(Man)-antigen administration indicators of development of regulatory immunity, including upregulation of *ikzf2*, which encodes Helios, although *foxp3* was not upregulated at this time point. Together, these T cell fates result in insufficient T cell help available to drive strong anti-OVA antibody responses.

**Figure 5:**
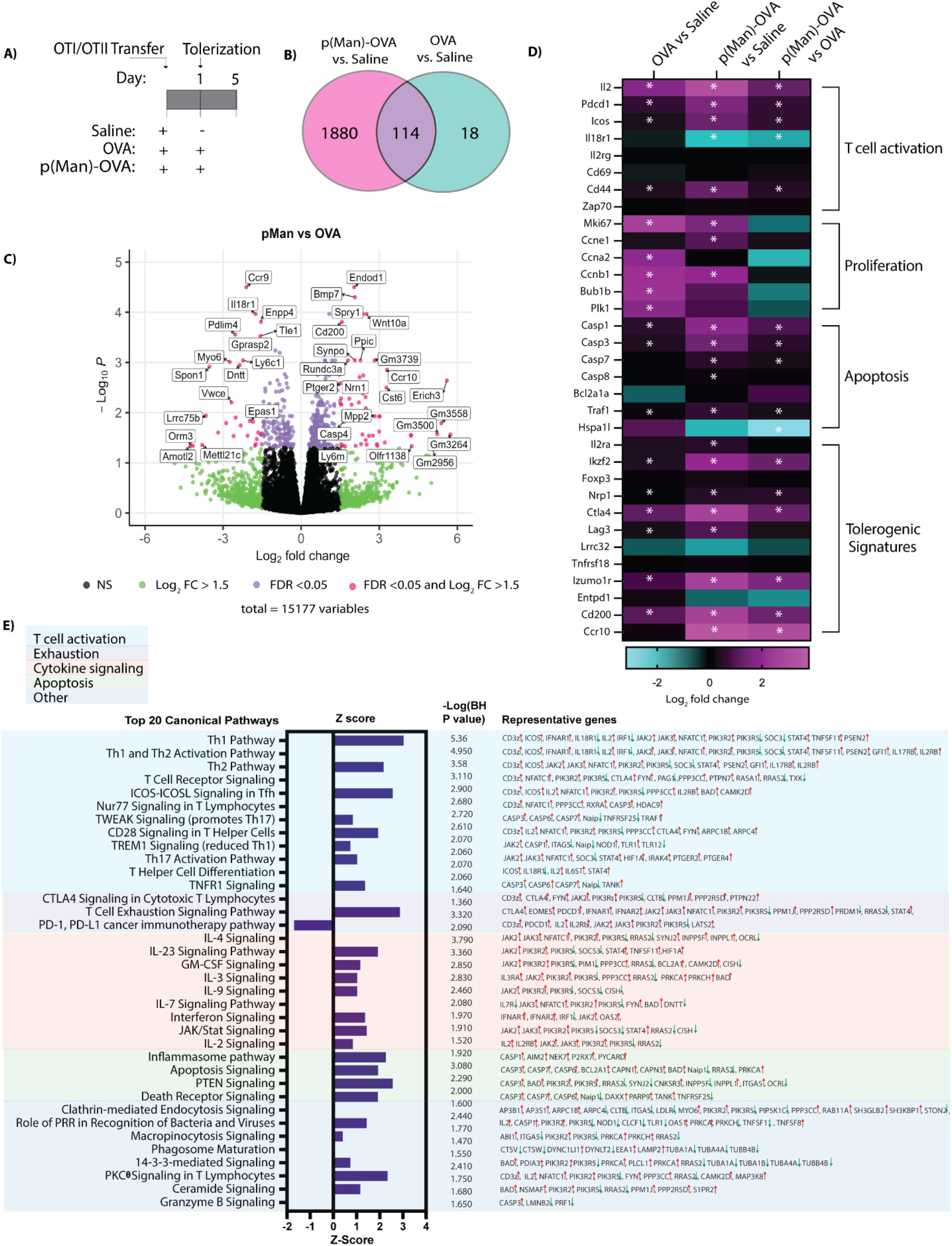
p(Man)-antigen treatment increases T cell intrinsic molecular signatures of TCR signaling along with apoptosis and exhaustion. (A) Strategy for RNA-seq. OT-I and OT-II T cells were adoptively transferred into C57Bl/6 mice on day 0. On day 1, mice were treated with p(Man)-OVA, free OVA or saline and OT-II cells were isolated from the spleens and lymph nodes for RNA-seq on day 5. *N=*4-5 samples per treatment group, 3 mice were pooled per sample. (B) Number of DEGs (FDR < 0.05; no FC cutoff) in p(Man)-OVA vs saline treated groups compared to OVA vs saline treated groups in OT-II. (C) Volcano plot of p(Man)-OVA vs OVA treatment in O-II. (D) Heatmap showing selected significant DEGs for the three comparisons (OVA vs saline; p(Man)-OVA vs saline; p(Man)-OVA vs OVA) of OT-II cells. Significant DEGs (FDR < 0.05; no FC cutoff) are indicated with an asterisk. (E) Significantly up-or down-regulated canonical immune pathways expressed in p(Man)-OVA groups compared with OVA groups as identified by ingenuity pathway analysis (IPA). Bars represent calculated z score. The BH-adjusted P value and associated genes are displayed to the right. Up/down arrows indicate gene expression.

## Discussion

In this study, we synthesized a novel mannosylated polymer, p(Man), that can be easily conjugated to any protein antigen (such as a biologic drug) to reduce the T follicular helper and resulting antibody responses against the unmodified antigen upon future administration. We showed that pre-treatment with p(Man)-antigen was able to significantly reduce the antigen-specific antibody response not only to OVA as a model protein but also to therapeutic doses of uricase and asparaginase, both highly immunogenic protein drugs^32,33^. We first demonstrated that p(Man) conjugation significantly increased the delivery of the conjugated antigen to the liver to a greater extent than even our previously published liver-targeted glycopolymers^22,23^. This difference may be due to the more promiscuous binding of mannose residues to a wider variety of C-type lectins compared to GalNAc or GlcNAc, resulting in increased endocytosis and antigen presentation^25,34–36^, or it may be mediated by signaling through a specific mannose-binding lectin in the liver we have yet to identify^37,38^. Our lab has previously shown that polymeric mannose can increase antigen uptake and presentation by a variety of APCs and this increased antigen presentation in the absence of co-stimulation may be a driving mechanism of tolerance in this system^26–28,39^. Like the previously published glycopolymers, pre-treatment with p(Man)-OVA resulted in an impaired CD4^+^ antigen-specific T cell response in the OT vaccination model, with an exceptional decrease in the number of Tfh cells within the OT-II population. Once we saw this lack of Tfh cell help, we confirmed that p(Man)-OVA pretreatment was sufficient to prevent an OVA-specific antibody response. p(Man)-OVA treatment fully prevented a detectable OVA-specific antibody response, while treatment with unmodified OVA, which contains native mannosylation^40^, did not. This provides evidence for the unique capacity of our linear p(Man) polymer to reduce humoral immune responses to an antigen over other types of mannosylation, at least native mannosylation in the chicken. To investigate the capacity for p(Man)-antigen treatment to prevent antibody responses to a highly immunogenic therapeutic protein, we conjugated it to uricase. Uricase is a tetrameric enzyme that degrades uric acid into the more easily excreted allantoin and is used as a therapeutic to treat hyperuricemia^41,42^. However, even when PEGylated, it elicits an immune response in the majority of patients, limiting therapeutic efficacy^43–45^. When p(Man)-uricase was given as a pre-treatment prior to multiple doses of unconjugated therapeutic uricase doses, we found a significant reduction in the uricase-specific immune response via a reduction in uricase-specific antibodies, fewer uricase-specific antibody-secreting cells, and fewer uricase-specific B cells in the spleen, indicating a broad reduction in the humoral response to this immunogenic antigen.

The effect of p(Man)-uricase treatment was different than p(Man)-OVA or p(Man)-asparaginase, as the antibody response to uricase was not fully eliminated. Despite the presence of detectible antibodies after p(Man)-uricase treatment, they were significantly reduced compared to untreated mice. This reduction may be clinically relevant, as the magnitude of the ADA response has been shown to affect the extent of clinical consequences^46^. Furthermore ADA-reducing therapeutics have demonstrated in both preclinical and clinical studies that a reduction in ADAs to uricase is correlated with reduced symptoms of gout such as serum uric acid control and reduction in tophi^47,48^. It is not yet clear how antigenic differences alter the efficacy of p(Man)-antigen treatment. The increased immunogenicity of uricase may be the result of its enzymatic function, which produces hydrogen peroxide, a reactive oxygen species, as a by-product, which may increase inflammation^49,50,51^. Alternatively, there may be differences in post-translational glycosylation of the different antigens, which has been shown to alter immune response^52,53^. The efficacy of p(Man)-antigen treatment may also differ across antigens due to the differences between the natural affinity of an individual’s T cell or B cell compartment to each antigen’s epitopes^12,54,55^.

To probe the immune mechanisms responsible for the effects of p(Man)-antigen delivery we performed an αCD25 depletion after p(Man) treatment. We found that pretreatment with p(Man)-uricase lead to a significant reduction in the uricase-specific antibody response despite the depletion. αCD25 depletion has been shown to deplete Tregs in addition to other recently activated CD25 expressing cells^56^. We interpret these data to suggest that Tregs generated during p(Man)-antigen treatment likely do not play a predominant role actively suppressing Tfh or B cells to cause the reduction of antibodies we see after p(Man)-antigen treatment. It is possible that the reduction of other CD25-expressing cells such as NK cells, B cells and activated T cells could confound this readout, but we do not expect the activated T cells or NK cells to have a role in maintaining the antibody response once germinal center activity has been initiated, and if critical antibody-generating B cells were depleted we would have expected antibody responses to decrease after treatment with αCD25. We also recognized the possibility that the presence of p(Man), a non-protein component and potential antigen, associated with the protein antigen during initial priming of B cells could affect downstream antibody responses though hapten immunodominance, resulting in a p(Man)-dominated antibody response. This immunodominance has been shown with 4-hydroxy-3-nitrophenylacetyl (NP)-OVA when more copies of NP are conjugated to an OVA protein, the B cells that are positively selected by T cells in the germinal center predominantly bind NP while expressing the OVA epitopes on their MHC after they endocytose the entire construct^57^. To test for this effect, we looked for antibody responses against the p(Man) polymer, but we saw no detectable response. Therefore, hapten immunodominance is unlikely to account for the reduction in antibody response we see to uricase after p(Man)-uricase treatment.

To further investigate mechanism, we performed transcriptional analysis on the OT-II T cells early (5 days) after p(Man)-OVA treatment and found upregulation of signatures associated with TCR-signaling, proliferation, apoptosis, and alternative mechanisms of T cell activation. These data suggest that p(Man)-antigen treatment increases presentation of the antigen to the T cells and resultant TCR signaling; however these T cells undergo non-Tfh fates of apoptosis, regulatory responses, exhaustion, or other non-germinal center responses limiting the Tfh help available to the B cells, ultimately preventing strong antibody responses against the antigen. Elevation in *nrp1*/*ikzf2*, genes that have canonically been used to differentiate thymic-derived Tregs from the FoxP3^+^ Tregs that are induced later in the periphery, show evidence of some of the earliest regulatory response to the antigen on our brief experimental timeline, even as the traditional CD25^+^FoxP3^+^ Tregs demonstrated to mediate effects in longer timeline depletion experiments may not have developed^58^. Of additional interest might be the elevation in chemokine receptor CCR10, which was found to be expressed by a subset of FoxP3^+^ Tregs homing via CCL28 secreted by cholangiocytes in the inflamed liver, where we hypothesize many of the lectin receptors capable of binding p(Man) for antigen uptake are located^31^. Lacking in costimulation from APCs that interacted with p(Man)-OVA, markers of anergy like *izumo1r* (FR4), as well as co-inhibitory receptors like *cd200, lag3,* and *ctla4* were also upregulated on OT-IIs in the pMan treated groups and likely represent key checkpoints that prevent ADA formation. Within the pathway analysis, we also saw upregulation of pathways associated with uptake such as endocytosis. This may be due to receptor endocytosis such as TCR and other surface receptor down-regulation associated with tolerance^59–61^ which could also account for some reduced T cell responses we see in the OT model.

Here we investigated the capacity of p(Man)-antigen treatment to prevent T cell dependent IgG antibody responses to subsequent antigen exposure in an antigen-specific manner. This technology has the potential to be applied in combination as a pretreatment to a number of immunogenic biologics in which ADA responses can limit therapeutic efficacy. Examples of potential use cases include preventing ADAs to chronically administered monoclonal antibody therapies such as adalimumab^62^, enzyme replacement therapies such as phenylalanine ammonia lyase for the treatment of phenylketonuria^63^ and factor VIII replacement for hemophilia A^64^.

With further validation, it may be used in combination with adeno-associated viral vectors to prevent ADAs and allow re-dosing^65^. In other studies, Cao et al. found that p(Man)-antigen pretreatment can be used to prevent food allergic responses and reduce antigen-specific IgE as well as IgG^66^. Future work can inform how p(Man) conjugation may impact IgE responses relevant in a minority of ADA responses^67,68^ and other diseases such as asthma^69^. Since we hypothesize p(Man)-antigen acts through the abrogation of T cell help, it may not be able to reduce T cell independent antibody responses, and further work should be done to investigate this application.

The experiments conducted in this study were all done in mouse models, however recent success of conceptually similar tolerance strategies have progressed to clinical trials. A glycosylation-modified antigen therapy has recently demonstrated safety and tolerability in a phase 1 clinical trial for celiac disease^70^ and is currently undergoing trials for multiple sclerosis (NCT04602390) as well. It is important to consider that while mannose-binding c-type lectins such as the mannose receptor have been identified as one-to-one orthologs between mice and humans, others have not^24^. Subsequent studies of p(Man)-antigen’s effects on antibody responses in humanized mice and non-human primates would further inform its translational potential.

In conclusion we have demonstrated that p(Man)-antigen conjugate therapy can significantly diminish the resulting antibody response to subsequent administration of an immunogenic antigen. This technology has the potential to be used with any immunogenic protein drug to prevent a patient from eliciting a strong ADA response to the drug and reduce the risk of adverse effects associated with ADAs, including loss of therapeutic efficacy, infusion reactions and anaphylaxis. The ability the reduce antibodies in an antigen-specific manner may have other broad applications in the treatment of antibody-mediated diseases such as asthma, allergy and antibody-mediated autoimmune diseases where specific antigens can be identified such as pemphigus vulgaris and myasthenia gravis ^71,72^. Further work is necessary to investigate how p(Man)-antigen therapy would function in the therapeutic setting after an antibody response has already developed. It could potentially be combined with antibody reducing treatments such as anti-CD20 treatments, plasma cell depletion or IVIG to reduce an antibody response and use p(Man)-antigen therapy to keep the antigen-specific response of interest low once broad immunosuppressants are discontinued. Ultimately, this modular platform may enhance tolerance to current and future biologics, enabling long-term therapeutic solutions for an ever-increasing healthcare problem.

## Acknowledgements

We thank the Cytometry and Antibody Core Facility at the University of Chicago. We thank The University of Chicago HIM Facility (RRID:SCR_017916). We thank the Center for Research Informatics, which is funded by the Biological Sciences Division at the University of Chicago with addition funding from the Institute for Translational Medicine, CTSA grant number UL1 TR000430 from the NIH, and the University of Chicago Comprehensive Cancer Center Support Grant (NIH P30CA014599). We thank Suzana Gomes for her laboratory support throughout this project.

## Author Contributions

Conceptualization, R.P.W, K.B., D.S.W., and J.A.H.; Methodology, R.P.W., A.C.T., K.B., and D.S.W.; Investigation, R.P.W., A.C.T., K.B., J.T.A., A.T.A., M.N., A.S., E.A.W., A.J.S., A.L.L., K.C.R., and D.S.W.; Visualization, R.P.W., and K.C.R.; Resources, M.R.R., A.J.S., and D.S.W.; Writing-Original Draft, R.P.W. and J.A.H.; Writing-Reviewing & Editing, R.P.W., K.C.R., A.T.A., D.S.W., and J.A.H.,; Funding Acquisition, J.A.H.; Supervision, D.S.W., and J.A.H.

## Data and materials availability

Gene expression data obtained by RNA-seq will be deposited in the NCBI Gene Expression Omnibus. All other data needed to evaluate the conclusions in the paper are present in the paper or the Supplementary Materials. The datasets generated and analyzed during the study are available from the corresponding author upon request.

## Methods

### Study Design

The purposes of this study were to test the feasibility of using p(Man)-antigen therapy to induce humoral tolerance and lessen an antigen-specific antibody response when administered before treatment with an immunogenic protein. We used OVA as a model protein along with TCR transgenic OT-I and OT-II to study antigen-specific T cell phenotypes and responses using flow cytometry and RNA-seq readouts. For RNA-seq experiments, *n=*4-5 samples were analyzed per treatment group, 3 mice were pooled per sample. For other OVA experiments, *n = 5* mice were used per group. We next tested p(Man)-antigen pre-treatment to reduce antibodies against immunogenic therapeutic proteins uricase from *Candida* and asparaginase from *E. coli.* Pre-immune mice with an initial antigen-specific antibody titer >2 prior to antigen experience were excluded from the study. *N = 8* mice were used per group in immunogenic challenge experiments. Each experiment was repeated at least twice.

### Mice

Mice were maintained in a specific pathogen-free facility at the University of Chicago. The experiments in this study were performed in accordance with the Institutional Animal Care and Use Committee. For OT-I and OT-II adoptive transfer experiments, 9 week old C57BL/6 mice were purchased from The Jackson Laboratory. OT-I (stock no: 003831) and OT-II (stock no: 004194) were crossed to CD45.1 mice (stock no: 002014) to yield congenically labeled OT-I and OT-II. For the immunogenic protein challenge experiments 9 week old female Balb/c mice were purchased from Charles River Laboratory.

### Transgenic T cell adoptive transfer

Spleens and lymph nodes (axillary, brachial, inguinal, and popliteal) were isolated from TCR transgenic mice. OT-I and OT-II T cells were isolated by negative magnetic bead selection using a CD8 and CD4 (Stemcell) negative selection kit, respectively. Cell purity was assessed by flow cytometry. OT-I and OT-II cells in DMEM were injected through the tail vein.

### Immunogenic antigens

All OVA injections were performed with 15 μg Endofit OVA (Sigma). All asparaginase injections were performed with 15 μg clinical wild-type asparaginase from *E. coli* (Asparaginase 5000, Medac GmbH). Asparaginase was purified via HIS-tag affinity purification and size exclusion chromatography. All uricase injections were performed with 0.5U *S. candida* uricase expressed in *E. coli* (Sigma) measured via Amplex red uricase activity assay (Thermo Fisher). *S. candida* uricase was purchased from Sigma and purified via FPLC on the Äkta system (Cytiva). Uricase was reconstituted in 50 mM sodium phosphate pH 6.4 and purified via anion exchange chromatography on a Sepharose Q FF column (Cytiva) followed by buffer exchange into 50 mM sodium carbonate bicarbonate pH 9.2 and size exclusion chromatography via HiLoad^TM^ 16/600 Superdex^TM^ 200pg (Cytiva). Activity of purified uricases and p(Man)-uricase constructs were measured using Amplex^TM^ Red Uric Acid/Uricase Assay kit (Thermo Fisher). Injections were administered once per week intravenously in the tail vein.

### p(Man) polymer synthesis

Unless otherwise stated, chemicals were reagent grade and purchased from Sigma-Aldrich. All NMR spectra were collected on a Bruker Avance-II 400 MHz NMR and analyzed with MnovaNMR (Mestrelab). GPC characterization was performed on Tosoh EcoSEC size exclusion chromatography System using Tosoh SuperAW3000 column at 50°C, with the eluent DMF with 0.01 M LiBr. A refractive index (RI) detector was used to detect polymers, and mass values were determined by column calibration with PMMA standards. Mannose monomer and azide-terminated RAFT chain transfer agent were synthesized as previously described^26,28^. Full synthesis schema with structures are provided in supplementary information.

For RAFT polymerization, mannosylated monomer (300 mg, 1.027 mmol) and HEMA (Combi Blocks, 300 mg, 2.326 mmol) were dissolved in dry DMF in a Schlenk tube. To that solution, azide CTA (17 mg, 0.029 mmol) and free radical initiator AIBN (1 mg, 0.0061 mmol) were added. The reaction vessel was degassed via four freeze-pump-thaw cycles and then heated to 70°C to initiate polymerization. After 14 hr, the reaction vessel was immersed in liquid nitrogen to stop the reaction. The polymer was the precipitated in cold diethyl ether three times. The final product was dried under reduced pressure. The product was white powder (550 mg, yield 89%). The polymer was characterized by ^1^H-NMR and GPC (**Supplemental Figures 1-3**) Resulting polymer had a number average molecular weight of 18 kDa.

### p(Man) antigen constructs

Azido-terminated p(Man) polymer was conjugated to lysine residues on the antigen via click chemistry using a self-immolative heterobifunctional linker, as previously described^22^. Conjugation was verified through an increase in molecular weight visible on SDS-PAGE separation (BioRad) which was reduced upon addition of 0.05% β-mercaptoethanol in the loading buffer. Mice were treated weekly for 2-3 weeks with 2.5-5μg p(Man)-antigen intravenously in the tail vein.

### Antigen biodistribution

C57BL/6 mice were treated by tail-vein injection with fluorescently labelled OVA in the form of free OVA (OVA_647_),OVA_647_–p(GalNAc), OVA_647_–p(GlcNAc); the subscripts indicate the fluorophores Dy-647 from Dyomics. After 3 hr, the livers of mice treated with OVA_647_, OVA_647_– p(GalNAc)or OVA_647_–p(GlcNAc) were perfused via the hepatic portal vein with 5–10 ml of 42°C Krebs–Ringer modified buffer (KRB) supplemented with 0.5 mM EDTA. The perfused livers and spleens were collected, placed in 4°C PBS, then analyzed for total fluorescence using an IVIS Spectrum in vivo imaging system (PerkinElmer).

### Preparation of single-cell suspensions

Draining lymph nodes (inguinal, popliteal, inguinal, axillary) and/or spleens were collected and filtered through a 70 μm filter. Lymph nodes were mechanically disrupted and digested at 37°C for 45 min in collagenase D. Digested lymph nodes and spleens were processed into single-cell suspensions via mechanical disruption and passage through a sterile 70 μm strainer. Red blood cells in splenocyte suspensions were lysed in ACK lysing buffer (Gibco) for 5 min before quenching with 15 mL DMEM. The single cell suspensions were then resuspended in IMDM + 10% FBS + 1% pen/strep.

### Flow cytometry

For phenotypic analysis of cells, 1.5×10^6^ cells were stained in PBS with 1:200 CD16/CD32 Fc Block (Biolegend) and 1:500 Live/Dead fixable dye (ThermoFisher) at 4°C for 15 min. Cells were washed then stained with 1:200 surface antibodies and multimers at 4°C for 30 min. Cells were washed then fixed in 2% paraformaldehyde. For intracellular staining, Cytofix/Cytoperm (BD Biosciences) was used at 4°C for 20 min, and cells stained intracellularly in Perm Wash Buffer (Biolegend) with 1:200 antibodies at 4°C for 30 min. For transcription factor stain, FoxP3 Transcription Factor Kit (eBioscience) was used per manufactures protocol. For complete antibody panel see **Supplemental Table 1**. Representative gating found in **Supplemental Figure 5**.

### Antigen-specific multimer production

Uricase was biotinylated using EZ-link NHS-biotin (Thermo Scientific). Unreacted NHS-biotin was removed using Zeba spin desalting columns, 7kDa MWCO (Thermo Scientific). The extent of biotinylation was measured using the QuantTag Biotin Quantification kit (Vector Laboratories) to ensure 1:1 molar ratio of uricase and biotin. Biotinylated uricase was reacted for 20 min on ice with 4:1 molar ratio of biotin to streptavidin-conjugated PE or streptavidin-conjugated APC (Biolegend). Streptavidin-conjugated FITC (BioLegend) was reacted with excess free biotin to form a non-antigen-specific streptavidin probe as a control. Multimer formation was confirmed using SDS-PAGE gel. Cells were stained for flow cytometry with all three streptavidin probes at the same time as other fluorescent surface markers at a volumetric ratio of 1:200. The antigen-specific staining of B cells with multimer was verified by staining with and without 5M uricase on splenocytes from vaccinated and antigen-naive mice.

### Antigen-binding ELISA

For all experiments, blood was collected from mice at least 4 days after last treatment or challenge dose and the plasma was isolated. Plasma was assessed for antigen-specific IgG by ELISA. 96-well ELISA plates (Costar Highbind flat-bottom plates, Corning) were coated with 10 μg/mL antigen in carbonate buffer (50 mM sodium carbonate/sodium bicarbonate, pH 9.6) overnight at 4°C. The following day, plates were washed three times in PBS with 0.05% Tween 20 (PBS-T) and then blocked with 1x casein (Sigma) for 2 h at RT. Following blocking, wells were washed three times with PBS-T and further incubated with six 10-fold dilutions of plasma in 1x casein for 2 hr at RT. Wells were then washed five times with PBS-T and incubated for an additional 1 hr at RT with horseradish peroxide (HRP)-conjugated antibodies against mouse IgG, IgG1, IgG2b, IgG2c, or IgG3 (Southern Biotech). After five washes with PBS-T, bound antigen-specific Ig was detected with tetramethylbenzidine (TMB) substrate. Stop solution (3% H_2_SO_4_ + 1% HCl) was added after 18 min of TMB incubation at RT, and the OD was measured at 450 and 570 nm on an Epoch Microplate Spectrophotometer (BioTek). Background signal at 570 nm was subtracted from the OD at 450 nm. The area under the curve (AUC) of background-subtracted absorbance versus log-transformed dilution was then calculated. Titer was determined as the log-transformed dilution at which the background-subtracted absorbance was greater than 0.01.

### Uricase-binding IgG ELISpot assay

Uricase was inactivated by heating at 98°C for 5 min. ELISpot plates (Millipore IP Filter plate) were coated with 20 µg/mL heat-inactivated uricase in sterile PBS overnight at 4°C. Plates were then blocked using ELISpot Media (RPMI 1640, 1% glutamine, 10% fetal bovine serum, 1% penicillin-streptomycin) for 2 hr at 37°C. Splenocytes from vaccinated mice were seeded in triplicate at a starting concentration of 6.75×10^5^ cell/well and diluted seven times in 3-fold serial dilutions. Plates were incubated for 18 hr at 37°C and 5% CO_2_ after which the cells were washed five times in PBS. Wells were incubated with 100 µL IgG-biotin HU adsorbed (Southern Biotech) for 2 hr at RT. Next, plates were washed four times in PBS followed by the addition of 100 µL HRP-conjugated streptavidin/well for 1 hr at RT. Plates were washed again and incubated with 100 µL TMB/well for 5 min. Finally, plates were washed three times with distilled water and left to dry completely in a laminar flow hood. A CTL ImmunoSpot Analyzer was used to image plates, count spots, and perform quality control.

### RNAseq Data Collection

Mice were adoptively transferred with 750,000 OT-I and OT-II cells at day 0. At day 1, mice received i.v. saline, 5 µg p(Man)-OVA, or equimolar OVA. Four days later, mice were sacrificed and spleens and liver and peripheral lymph nodes were pooled. OT-II cells were sorted into Trizol using fluorescence activated cell sorting (FACS) and frozen at −80°C. RNA was extracted (Qiagen, RNeasy Micro), and three mice were pooled per replicate. RNA integrity was measured by the Agilent 2100 Bioanalyzer. RNA was converted to cDNA using SmartSeq-v4 (Takara, cat no. R400752). The library was prepared using Nextera XT DNA library preparation kit (Illumina, cat no. FC-131-1096). For each setting, all samples were pooled and sequenced using Illumina HiSeq 4000 (2×100 paired end).

### RNAseq analysis

The RNA-seq analysis was performed under the R programming and software environment for statistical computing and graphics version 3.6 (R Core Team, 2019). We assessed the quality of FastQ files using the FastQC tool (version 0.11.5). Raw reads were aligned to the GRCh38 primary genome assembly using Spliced Transcripts Alignment to a Reference (STAR) aligner (version 2.7.2a) 1-pass algorithm^73^. Bam files were sorted in lexicographical order with the sambamba program (v0.5.4). Reads were assigned to exon features annotated in Ensembl Mus musculus GRCm38 annotation (release 97) using the FeatureCounts tool from the subread package (version 1.5.2) and the read counts were summarized by genes^74,75^. Picard tools (version 2.18.7) was used for post alignment quality control. Alignment-free transcriptome quantification method kallisto (v0.46.1) was used to estimate the transcript abundance of each sample^76^. Transcript-level estimates were summarized for gene-level analysis using R package tximport. Genes with zero read counts across all samples were removed. Raw library sizes were scaled using normalization factors calculated with the calcNormFactors function in the edgeR R package with trimmed mean of M-values (TMM) option enabled^77^. The normalized count per million (CPM) values were log_2_-transformed. We used the voomWithQualityWeights function from the limma package to remove heteroscedascity from the count data. A linear model for each gene was fit with the limma algorithm, adjusted for any batch effects, and the genes were ranked by differential expression using the empirical Bayes method. Differentially expressed genes were identified with the Benjamini-Hochberg procedure for multiple testing correction. Significance was set at an adjusted P-value threshold of 0.05. We used the Ingenuity Pathway Analysis Fall 2020 release (Qiagen) to identify canonical pathways, causal networks, and upstream regulators that related to significant differentially expressed genes in the p(Man)-OVA to OVA comparisons.

### CD25 depletion study

Mice were treated weekly for 3 weeks with either p(Man)-uricase or saline. Two weeks following last treatment, mice were intravenously injected with 500 μg of either anti-CD25 depletion antibody of the IgG2a isotype control (BioXCell). Two weeks following depletion mice were challenged weekly intravenously with 0.5U uricase for 4 wk.

### T cell depletion study

Mice were treated weekly for 3 wk with 500 μg αCD4 and αCD8 depleting antibodies or IgG2b isotype control antibodies (BioXCell). 4 days after initial depletion mice were administered 1U of uricase or saline i.v. weekly for 3 wk. Serum was collected for antibody measurement.

### Statistical Analysis

Statistically significant differences between experimental groups were determined using Prism software (v9, GraphPad). All statistical analyses are stated specifically in the figure legends for all experiments. For most experiments, unless otherwise specified in figure legend, one-way ANOVA was performed with a Tukey’s test to correct for multiple comparisons. For showing statistical significance ***P≤0.001; **P≤0.01; *P≤0.05, unless otherwise stated. For the RNA-seq experiment, a detailed description of the statistical analysis can be found in Methods. Analyses were performed using R and GraphPad Prism V.7 software. The threshold for statistical significance was fixed at p < 0.05.

