## Supplemental Materials for "Synthetically mannosylated antigens induce antigen-specific humoral tolerance and reduce anti-drug antibody responses to immunogenic biologics"

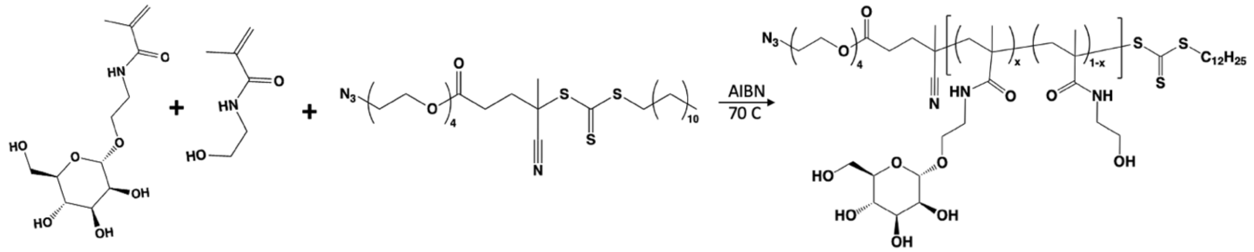


**Fig S1. Schema of p(Man) polymerization**

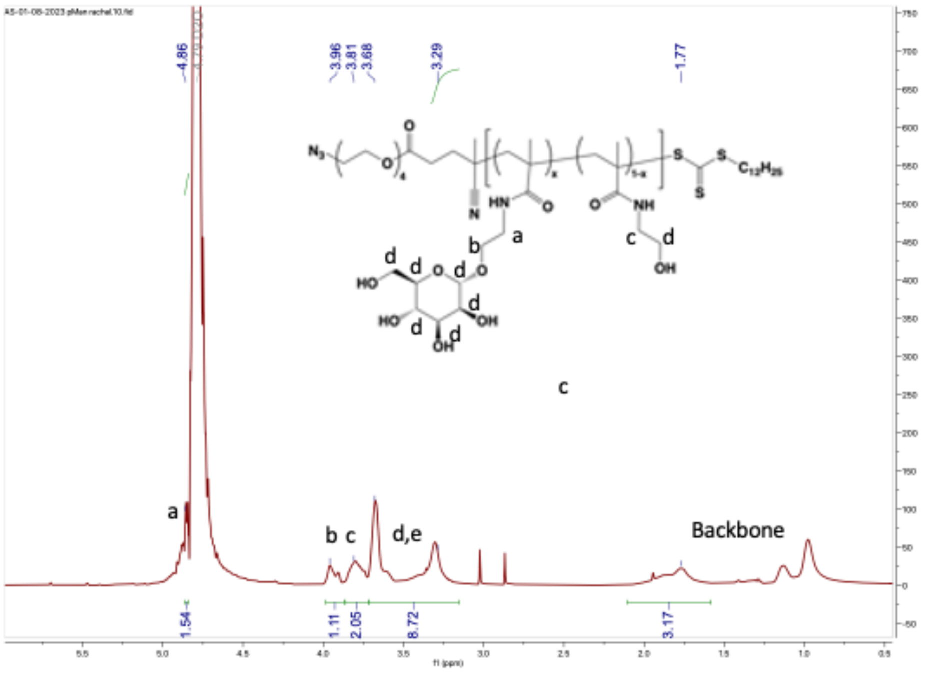


**Fig. S2: p (Man) NMR**

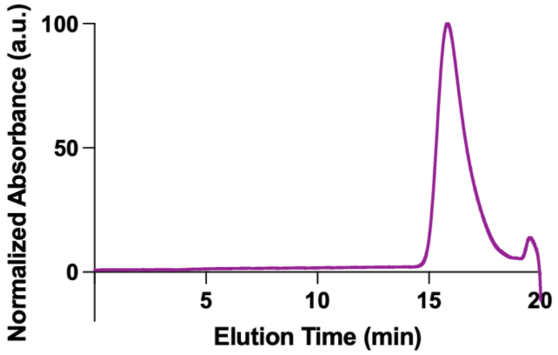

**Fig. S3: p(Man) GPC**

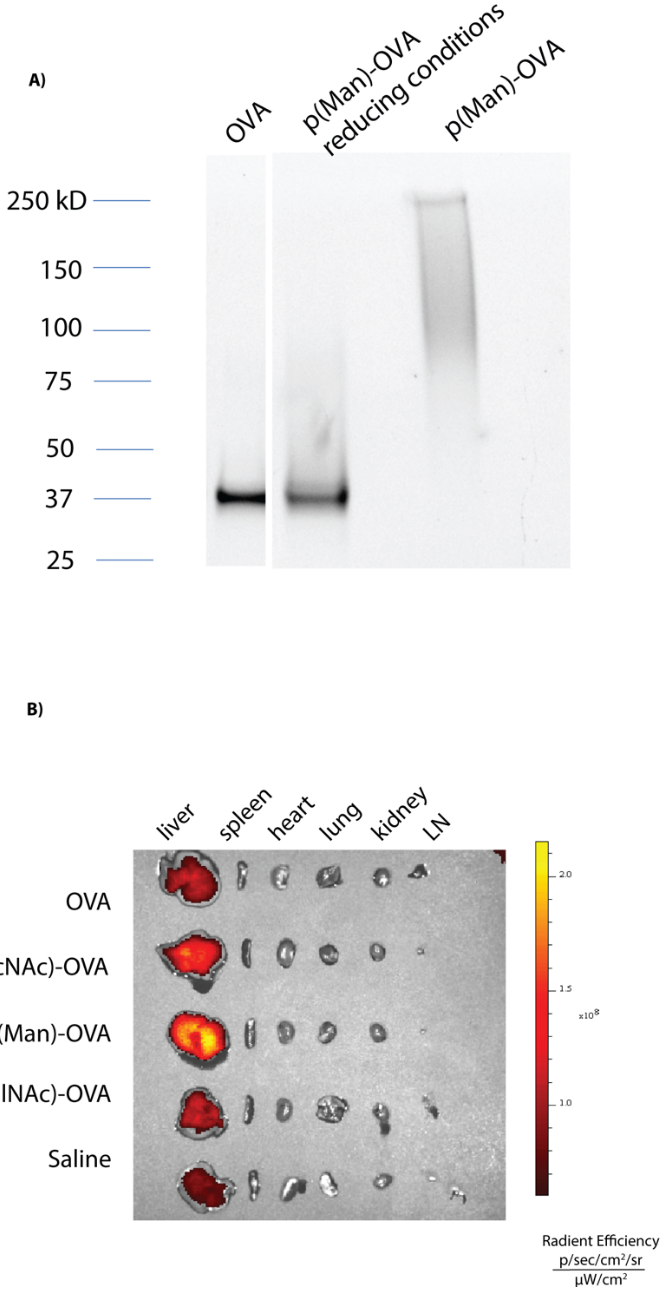


**Fig. S4: p(Man)-OVA characterization and biodistribution** (A) SDS-PAGE of unmodified OVA and p(Man)-OVA with and without β-mercaptoethanol to show self-immolative linker. (B) Representative IVIS image of liver, spleen, heart, lung, kidney and lymph nodes 3 hours after injection with OVA$_{AF647,}$ p(GlcNAc)-OVA$_{AF647,}$ p(Man)-OVA$_{AF647,}$ p(GalNAc)-OVA$_{AF647,}$ or saline iv.

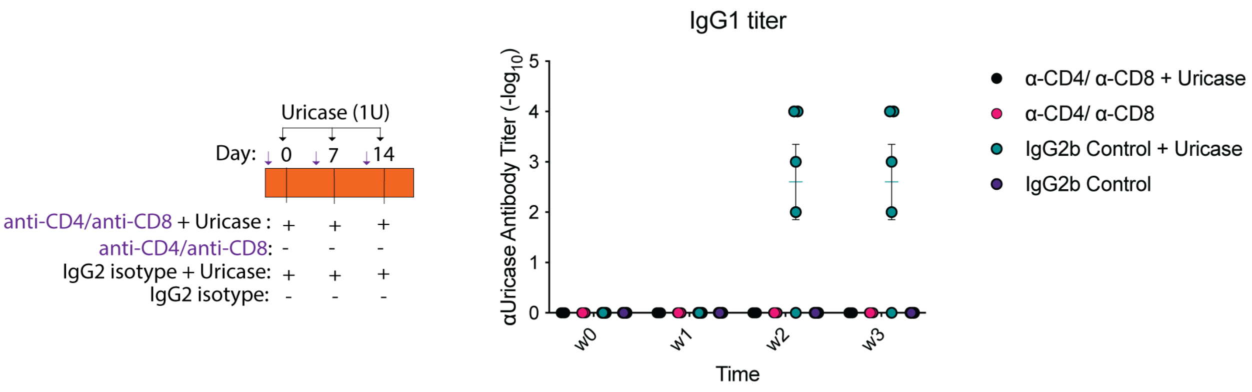


**Fig. S5: *Candida* uricase elicits a T cell-dependent antibody response in mice** (A) *N*=4 Balb/c mice were treated weekly iv with 1U of *Candida* uricase along with αCD4 and αCD8 depletion antibodies or an equivalent of IgG2b isotype control. (B) Time-course of uricase-specific IgG antibody response represented as $\log_{10}$ titer. Symbols represent individual mice.

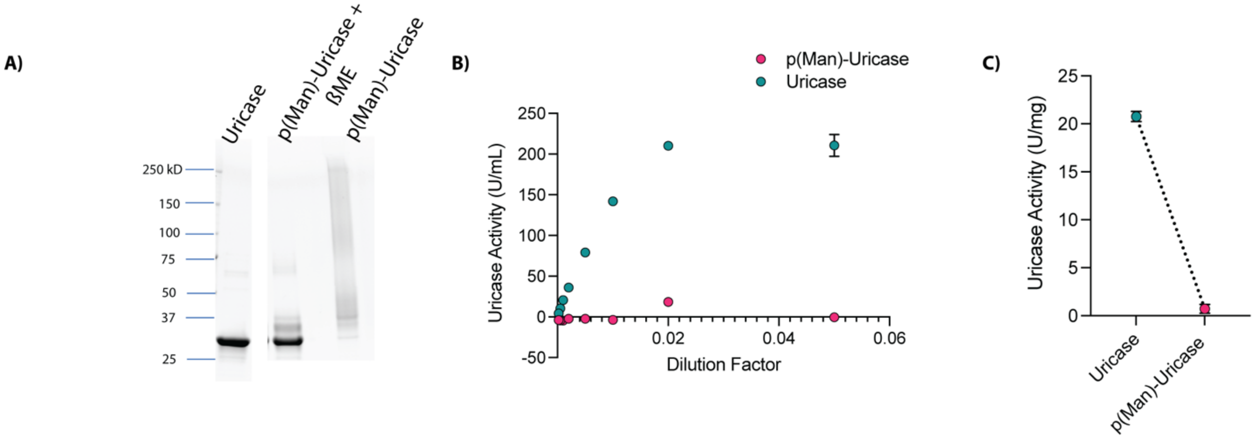


**Fig. S6: Characterization of p(Man)-uricase conjugates** (A) SDS-PAGE of *Candida* uricase compared to p(Man)-conjugated uricase with and without β-mercaptoethanol. (B) Uricase activity (U/mL) of p(Man)-conjugated and native uricase across dilutions. (C) Uricase activity (U/mg) of p(Man)-conjugated and native uricase.

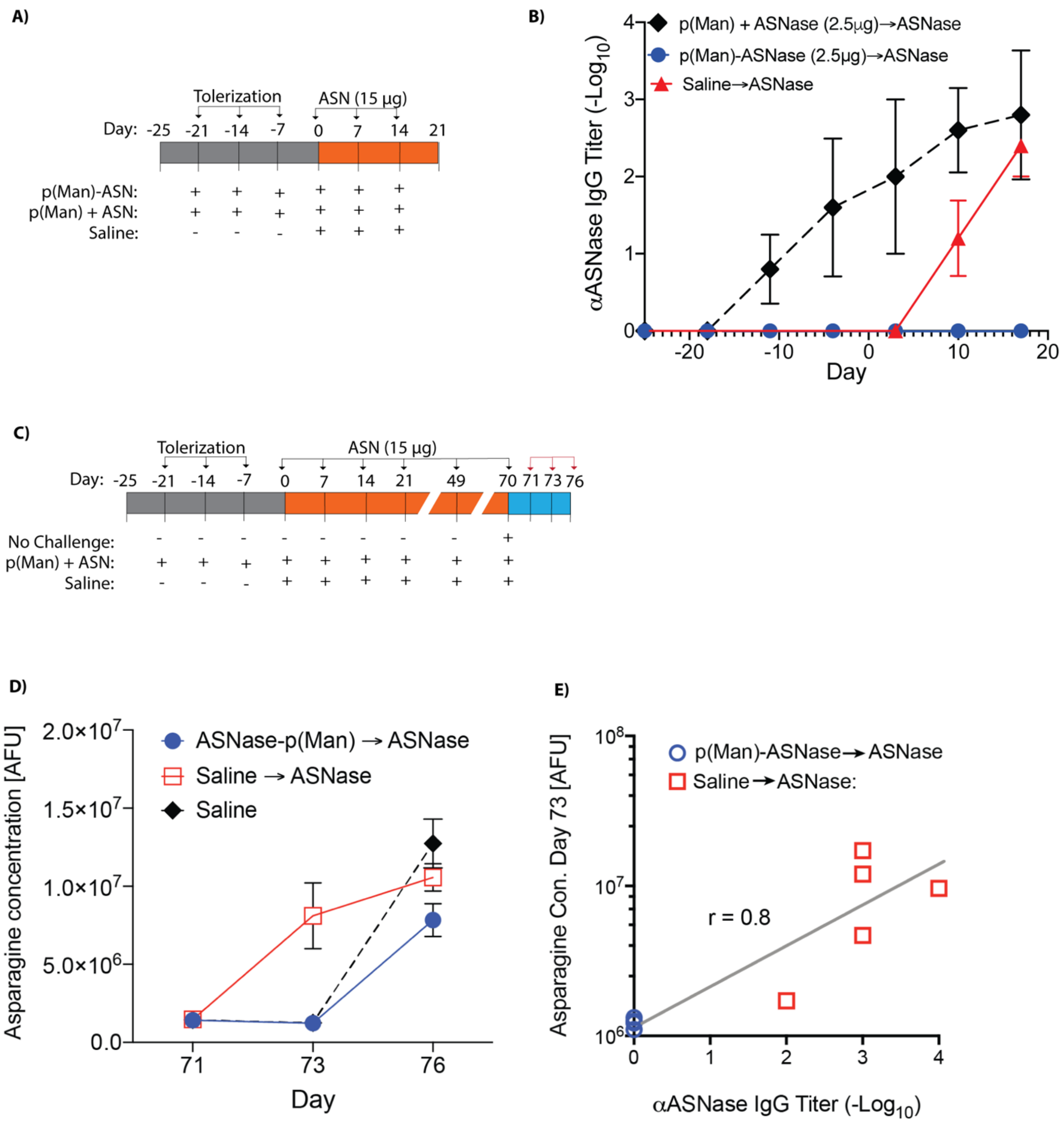


**Fig. S7: p(Man)-asparaginase pre-treatment prevents antibody response to *E Coli* asparaginase** (A) *N=5* Balb/c mice were treated weekly for 3 weeks with p(Man)-asparaginase followed by 3 weeks of challenge with high dose asparaginase. (B) Time-course of the asparaginase-specific IgG antibody response represented as $\log_{10}$ titer. (C) *N*=5 Balb/c mice were treated as in (A) followed by 4 weeks of challenge with high dose asparaginase after which mice were rested for 28 days before rechallenge on day 49 and 70. Serum was collected on days 71, 73, and 76 for measurement of asparagine concentration. (D) Serum asparagine concentration (AFU) of mice pre-treated with p(Man)-asparaginase or saline on days 71, 73, and 76. (E) Correlation between asparaginase-specific antibody response and asparagine concentration on day 73. Data is represented as means± SEM.

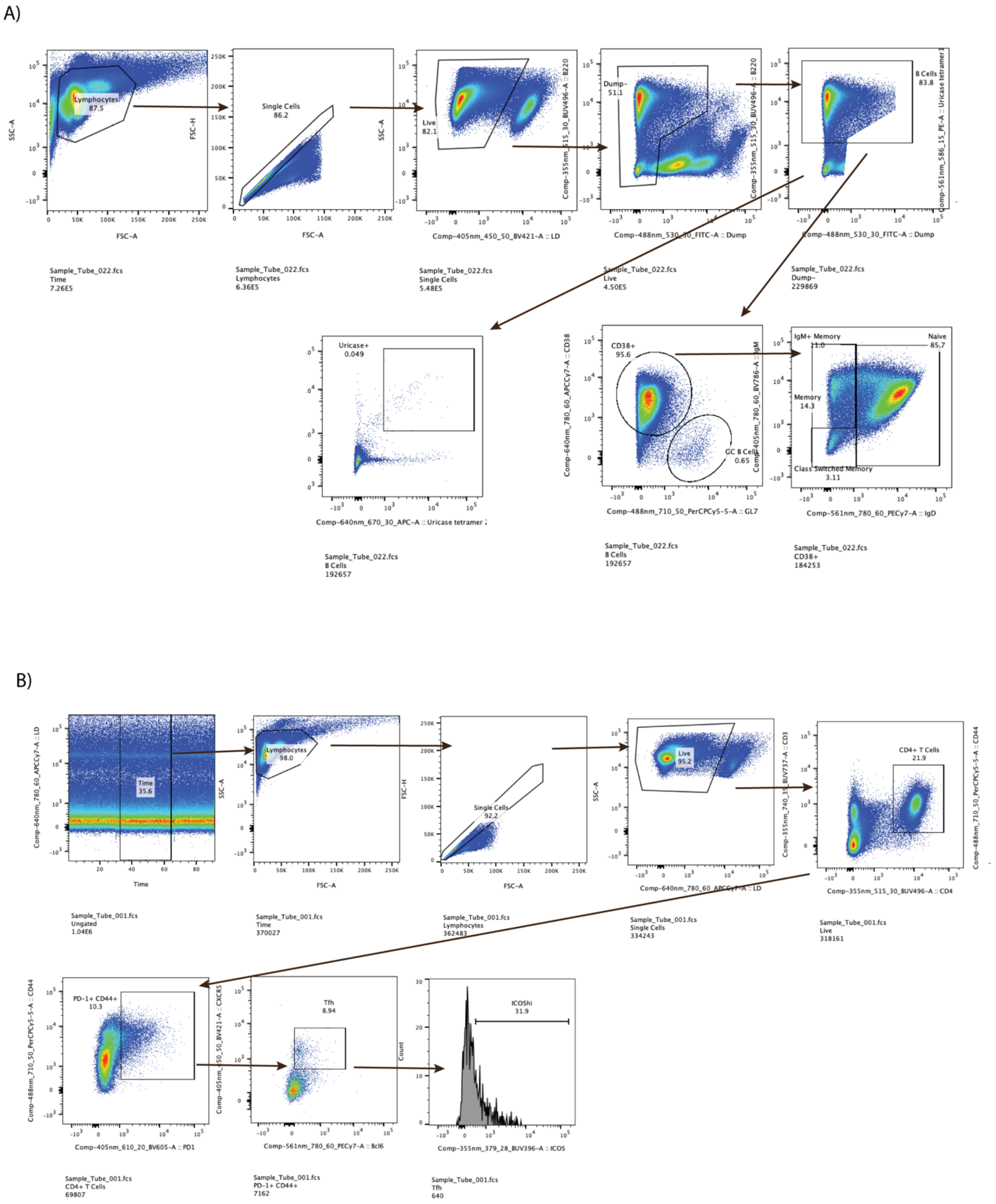

A)
Lymphocytes
87.5
SSC-A
FSC-A
Single Cells
86.2
FSC-H
Live
82.1
Comp-405nm_450_50_BV421-A :: LD
Dump-
51.1
Comp-355nm_515_30_BUV496-A :: B220
Comp-488nm_530_30_FITC-A :: Dump
B Cells
83.8
Comp-561nm_586_15_PE-A :: Uricase tetramer 1
Sample_Tube_022.fcs
Time
7.26E5
Sample_Tube_022.fcs
Lymphocytes
6.36E5
Sample_Tube_022.fcs
Single Cells
5.48E5
Sample_Tube_022.fcs
Live
4.50E5
Sample_Tube_022.fcs
Dump-
229869
Uricase+
0.049
Comp-640nm_670_30_APC-A :: Uricase tetramer 2
CD38+
95.6
GC B Cells
0.65
Comp-640nm_780_60_APCCy7-A :: CD38
Comp-488nm_710_50_PerCPCy5-5-A :: GL7
IgM+ Memory
11.0
Naive
85.7
Memory
14.3
Class Switched Memory
3.11
Comp-405nm_780_60_BV786-A :: IgM
Comp-561nm_780_60_PECy7-A :: IgD
Sample_Tube_022.fcs
B Cells
192657
Sample_Tube_022.fcs
B Cells
192657
Sample_Tube_022.fcs
CD38+
184253
B)
Time
35.6
Comp-640nm_780_60_APCCy7-A :: LD
Time
Lymphocytes
98.0
SSC-A
FSC-A
Single Cells
92.2
FSC-H
FSC-A
Live
95.2
SSC-A
Comp-640nm_780_60_APCCy7-A :: LD
CD4+ T Cells
21.9
Comp-355nm_740_35_BUV737-A :: CD3
Comp-355nm_515_30_BUV496-A :: CD4
Comp-488nm_710_50_PerCPCy5-5-A :: CD44
Sample_Tube_001.fcs
Ungated
1.04E6
Sample_Tube_001.fcs
Time
370027
Sample_Tube_001.fcs
Lymphocytes
362483
Sample_Tube_001.fcs
Single Cells
334243
Sample_Tube_001.fcs
Live
318161
PD-1+ CD44+
10.3
Comp-488nm_710_50_PerCPCy5-5-A :: CD44
Comp-405nm_610_20_BV605-A :: PD1
Tfh
8.94
Comp-405nm_450_50_BV421-A :: CXCR5
Comp-561nm_780_60_PECy7-A :: Bcl6
ICOShi
31.9
Count
Comp-355nm_379_28_BUV396-A :: ICOS
Sample_Tube_001.fcs
CD4+ T Cells
69807
Sample_Tube_001.fcs
PD-1+ CD44+
7162
Sample_Tube_001.fcs
Tfh
640

**Fig. S8: Flow cytometry gating strategy** (A) Gating strategy for the identification of Dump- B220+ B cell subsets and uricase-specific multimer-double-positive B cells. (B) Gating strategy for the identification of $CD3^+$ $CD4^+$ $CD44^+$ $PD1^+$ $CXCR5^+$ $Bcl6^+$ T follicular helper (Tfh) cells.

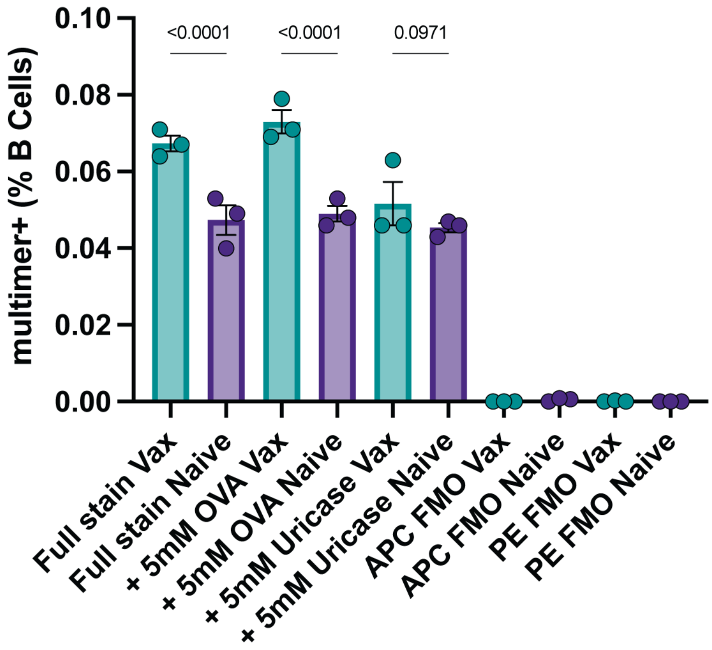


**Fig. S9: Validation of the antigen-specificity of the fluorescent uricase multimers** Splenocytes from naïve Balb/c mice or mice vaccinated with 0.5U iv uricase challenge were stained with APC-uricase and PE-uricase multimers with or without pre-incubation with 5mM OVA or uricase. Multimer-double-positive cells were quantified as a percentage of B220+ B cells. Data are shown as means ± SEM. Symbols represent individual mice and statistical differences in all graphs were determined by one-way ANOVA with Tukey's, p values are displayed on graph.

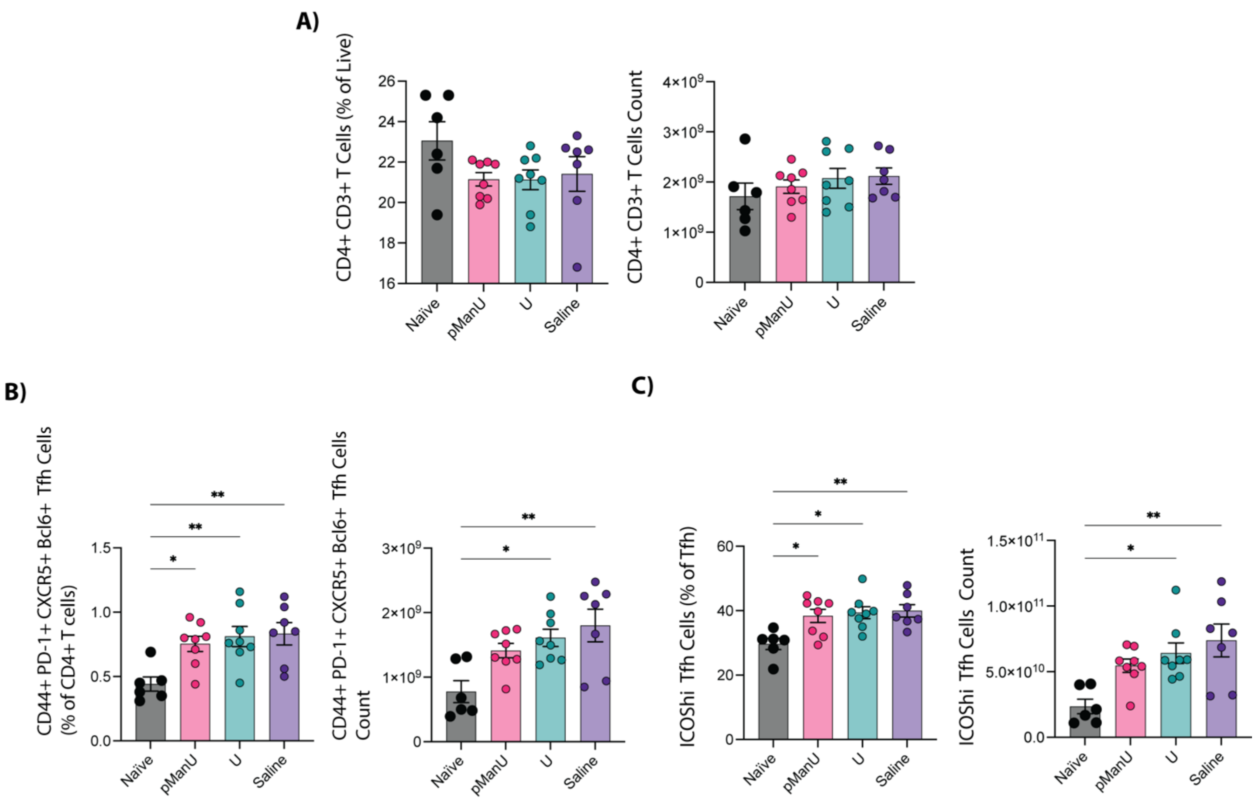


**Fig. S10: T cell compartment after p(Man)-uricase treatment** (A) Frequency of $CD3^+$ T cells as a percentage of live cells after pre-treatment with p(Man)-uricase, uricase or saline as in 2A. (B) Frequency of $CD4^+CXCR5^+PD\text{-}1^+Bcl6^+$ Tfh cells as a percentage of T cells after treatment. (C) Frequency of $ICOS^{hi}$ Tfh cells as a percentage of Tfh after treatment. Data are shown as means ± SEM. Unless otherwise stated symbols represent individual mice and statistical differences in all graphs were determined by one-way ANOVA with Tukey's *$p<0.05$, **$p<0.01$ and ***$p<0.001$.

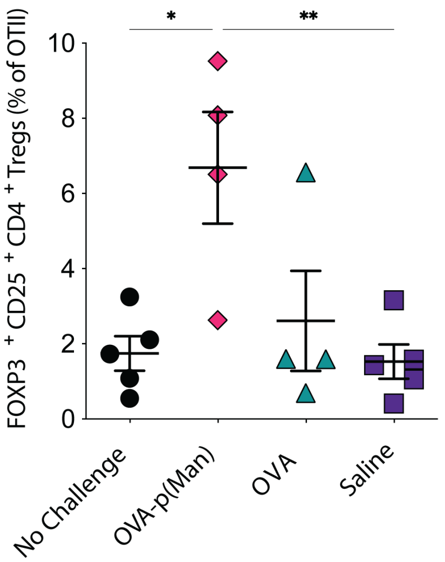


**Fig. S11: p(Man)-OVA treatment results in increased frequency of OVA-specific Tregs**

C57Bl/6 mice were treated as in Figure 1D. Frequency of OVA-specific $CD4^+$ $CD25^+$ $FoxP3^+$ T regulatory (Treg) cells were quantified as a percentage of OTIIs in the dLN. Data are shown as means ± SEM. Symbols represent individual mice and statistical differences in all graphs were determined by one-way ANOVA with Tukey's *$p<0.05$, **$p<0.01$ and ***$p<0.001$.

**Table S1. Flow cytometry antibody panels**

19 Day Adoptive Transfer Antibodies

| **Marker** | **Fluorophore** | **Vendor** | **Cat** |
|---|---|---|---|
| CD45.1 | BV421 | Biolegend | 110731 |
| CD8 | BV510 | BD | 563068 |
| CD4 | BV786 | BD | 563727 |
| CXCR5 | BV650 | Biolegend | 145517 |
| Bcl6 | PE-Cy7 | Biolegend | 358512 |
| Viability dye | e780 | Invitrogen | 65-0865-14 |
| CD3 | BUV395 | BD | 563585 |
| PD-1 | PerCP-Cy5.5 | Biolegend | 135208 |
| CD25 | PE | Biolegend | 101904 |
| FoxP3 | AF488 | BD | 560403 |

B Cell Panel Antibodies

| **Marker** | **Fluorophore** | **Vendor** | **Cat** |
|---|---|---|---|
| Viability dye | BV421 | Invitrogen | L34955 |
| CD11c | FITC | BD | 557400 |
| GR-1 | FITC | Biolegend | 108406 |

| | | | |
|---|---|---|---|
| CD4 | FITC | Biolegend | 100406 |
| CD8a | FITC | Biolegend | 100706 |
| B220 | BUV496 | BD Horizon | 612950 |
| CD19 | BUV395 | BD Horizon | 563557 |
| CD138 | BV605 | Biolegend | 142531 |
| IgM | BV786 | BD Optibuild | 743328 |
| CD38 | APC-Cy7 | Biolegend | 102728 |
| IgD | PE-Cy7 | Biolegend | 405720 |
| GL7 | PerCP-Cy5.5 | Biolegend | 144609 |

Tfh Cell Panel Antibodies

| **Marker** | **Fluorophore** | **Vendor** | **Cat** |
|---|---|---|---|
| Viability dye | e780 | Invitrogen | 65-0865-14 |
| ICOS | BUV395 | BD Horizon | 565885 |
| CD3 | BUV737 | BD Horizon | 612803 |
| CXCR5 | BV421 | Biolegend | 145512 |
| CD4 | BUV496 | BD Horizon | 612952 |
| CD44 | PerCP-Cy5.5 | BD Horizon | 563058 |
| PD-1 | BV605 | Biolegend | 135219 |
| Bcl6 | PE-Cy7 | Biolegend | 358512 |
| FoxP3 | AF647 | BD | 560402 |
| CD25 | PE | Biolegend | 101904 |
| CD44 | PerCP-Cy5.5 | Biolegend | 103032 |
